# *SLC1A5* provides glutamine and asparagine necessary for bone development in mice

**DOI:** 10.1101/2021.06.23.449651

**Authors:** Deepika Sharma, Yilin Yu, Leyao Shen, Guo-Fang Zhang, Courtney Karner

**Affiliations:** Department of Orthopaedic Surgery, Duke University School of Medicine, 300 North Duke Street, Durham, NC 27701, USA; Department of Internal Medicine, University of Texas Southwestern Medical Center, 300 North Duke Street, Durham, NC 27701, USA; Sarah W. Stedman Nutrition and Metabolism Center & Duke Molecular Physiology Institute, Duke University Medical Center, 300 North Duke Street, Durham, NC 27701, USA; Department of Medicine, Duke University School of Medicine, Durham NC 27701, USA; Charles and Jane Pak Center for Mineral Metabolism and Clinical Research. University of Texas Southwestern Medical Center at Dallas. Dallas, TX, 75390

## Abstract

Osteoblast differentiation is sequentially characterized by high rates of proliferation followed by increased protein and matrix synthesis, processes that require substantial amino acid acquisition and production. How osteoblasts obtain or maintain intracellular amino acid production is poorly understood. Here we identify *Slc1a5* as a critical amino acid transporter during bone development. Using a genetic and metabolomic approach, we show *Slc1a5* acts cell autonomously in osteoblasts to import glutamine and asparagine. Deleting *Slc1a5* or reducing either glutamine or asparagine availability prevents protein synthesis and osteoblast differentiation. Mechanistically, glutamine and asparagine metabolism support amino acid biosynthesis. Thus, osteoblasts depend on *Slc1a5* to provide glutamine and asparagine, which are subsequently used to produce non-essential amino acids and support osteoblast differentiation and bone development.

## Background

Osteoblasts are secretory cells responsible for producing and secreting the Collagen Type 1 rich extracellular bone matrix. Osteoblasts differentiate from mesenchymal progenitors in a well-coordinated temporal sequence regulated by the transcription factors RUNX2 and OSX (encoded by *Sp7*)^1,2^. Osteoblast progenitors are proliferative before undergoing terminal differentiation into postmitotic Collagen Type 1 (COL1A1) matrix producing osteoblasts^3–7^. Both proliferation and matrix production are biosynthetically demanding and burden osteoblasts with enhanced metabolic demands. For example, proliferation requires cells to increase nutrient and amino acid acquisition to generate the biomass necessary to duplicate cell mass for division. Likewise, matrix production places similar demands upon osteoblasts. Thus, bone formation is associated with increased synthetic demands due to osteoblast proliferation, differentiation and bone matrix production^8–11^.

These enhanced synthetic demands are predicted to require a constant supply of amino acids to sustain both proliferation and bone matrix production. Cells can obtain amino acids via uptake from the extracellular milieu or from *de novo* synthesis from glucose or other amino acids^10,12–16^. In osteoblasts, amino acid consumption is known to be transcriptionally regulated^17,18^ and rapidly increases during differentiation and in response to osteoinductive signals^19–23^. Importantly, genetic mutations that limit amino acid uptake are associated with decreased proliferative capacity and reduced bone formation^17,18,24^. Despite this, little is known about how osteoblasts obtain the amino acids necessary to promote robust proliferation and bone matrix synthesis.

The regulation of amino acid supply is an important regulatory node that is frequently upregulated to support proliferation and biosynthesis in many pathological conditions^25–28^. Amino acid uptake is controlled by a diverse array of membrane-bound transport proteins that transport amino acids into and out of the cell. Once inside the cell, amino acids have diverse fates. For example, amino acids may contribute directly to protein synthesis, facilitate signaling or be metabolized to generate ATP or other intermediate metabolites including other amino acids^29–32^. How osteoblasts obtain or utilize amino acids is not well understood. We recently identified Alanine, Serine, Cysteine transporter 2 (ASCT2, encoded by *Slc1a5)* as a potential regulator of amino acid supply in osteoblast progenitors^20,24^ ASCT2 is a Na^+^ dependent neutral amino acid exchanger that can transport glutamine, alanine, serine, asparagine and threonine^33–37^ In cancer cells, *Slc1a5* upregulation is associated with metabolic reprograming and is necessary for increased proliferation and biosynthesis^28,38–43^. In comparison, little is known about the role of *Slc1a5/ASCT2* dependent amino acid uptake during bone development.

Here we define the role of amino acid uptake through *Slc1a5/ASCT2* during bone development and homeostasis. Using a genetic and metabolic approach we demonstrate *Slc1a5* is required for osteoblast proliferation, differentiation and bone matrix production. Mechanistically, *Slc1a5* provides glutamine and asparagine to maintain intracellular amino acid homeostasis in osteoblasts. Collectively, these data highlight the previously unknown role for *Slc1a5/*ASCT2 in osteoblasts regulating differentiation and bone development.

## Results

### *Slc1a5* is required for bone development

We previously identified *Slc1a5* as a potential regulator of proliferation and WNT induced osteoblast differentiation in stromal cells^20,24^ To understand if *Slc1a5* functions in osteoblasts, we first characterized *Slc1a5* expression during osteoblast differentiation. *Slc1a5* is highly expressed in both naïve calvarial cells and bone marrow stromal cells and significantly increases during osteoblast differentiation (Fig. 1A). To determine if *Slc1a5* is required for osteoblast differentiation, we first utilized a CRISPR/Cas9 approach to knock out *Slc1a5* in cultured calvarial cells (Fig. S1A). Western blot analyses confirmed this approach effectively ablated ASCT2 protein (Fig. S1B). *Slc1a5* targeting did not affect early osteoblast differentiation but did prevent the induction of the mature osteoblast genes *Ibsp* and *Bglap* and prevented matrix mineralization in primary calvarial cells (Fig. 1B-C). To determine if *Slc1a5* dependent amino acid uptake is important for osteoblast differentiation *in vivo,* we generated a conditional (floxed) *Slc1a5* allele *(Slc1a5^fl^*) using homologous recombination (Fig. S1C). These mice were crossed with the *Sp7GFP:Cre* deleter mouse *(Sp7Cre)* that expresses GFP and Cre recombinase under the control of the *Sp7* promoter to generate mice lacking *Slc1a5* in committed osteoblast progenitors^44^. Western blot analysis confirmed the specific ablation of ASCT2 in bones isolated from *Sp7Cre;Slc1a5^fl/fl^* mice (Fig. S1D). *Sp7Cre;Slc1a5^fl/fl^* mice were characterized by delayed endochondral and intramembranous ossification evident at both embryonic (E) day E14.5 and E15.5 (Fig. 1D-E, H-I). In addition to delayed mineralization, *Sp7Cre;Slc1a5^fl/fl^* mice had reduced size of various skeletal elements (exemplified by the humerus, Fig. 1F and J) relative to *Sp7Cre;Slc1a5^fl/+^* littermate controls (Fig. 1F-G, J-K, S1E-L). By E16.5 neither the extent of mineralization nor the overall length of individual skeletal elements were significantly different from *Sp7Cre;Slc1a5^fl/+^* littermate controls suggesting there is a transient delay in endochondral ossification (Fig. 1L-O, S1M-X). Conversely, *Sp7Cre;Slc1a5^fl/fl^* mice are characterized by impaired intramembranous ossification at all stages evaluated (Fig. 1D-E,H-I,L-M,P-Q, S1Q-R). At birth, *Sp7Cre;Slc1a5^fl/fl^* calvariae had increased porosity (1.0±0.2 vs 2.3±0.2 in *Sp7Cre;Slc1a5^fl/+^* and *Sp7Cre;Slc1a5^fl/fl^* respectively, p<0.05), increased suture width (1.0±0.2mm vs 1.4±0.3 in *Sp7Cre;Slc1a5^fl/+^* and *Sp7Cre;Slc1a5^fl/fl^* respectively, p<0.05) and significantly less bone volume (0.7±0.02% vs 0.6±0.05% in *Sp7Cre;Slc1a5^fl/+^* and *Sp7Cre;Slc1a5^fl/fl^* respectively, p<0.05) as measured by micro-computed tomography (μCT) (Fig. 1P-S). Von Kossa staining of histological sections confirmed the reduction in mineralized bone matrix in newborn *Sp7Cre;Slc1a5^fl/fl^* calvariae (Fig. 1T-U). Thus, *Slc1a5* expression in osteoprogenitors is essential for bone development.

**Fig. 1.**
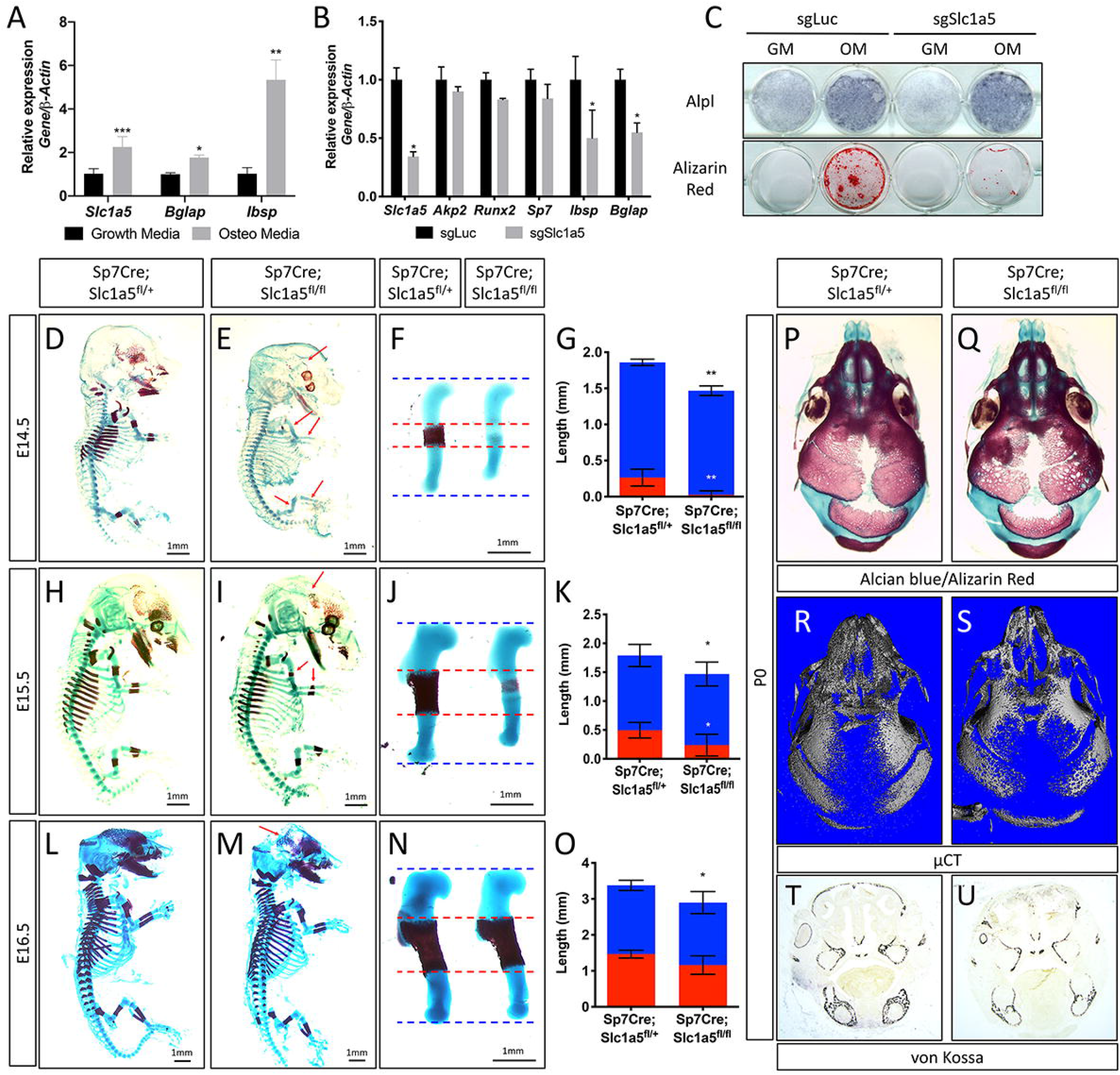
Slc1a5 is required for bone development in mice. **(A)** qRT-PCR analyses of gene expression in calvarial osteoblasts (cOB) cultured for 7 days in growth or osteogenic media. (**B-C**) qRT-PCR analyses **(B)** or functional assays **(C)** of the effect of *Slc1a5* deletion on osteoblast differentiation in cOB cultured for 7 days in osteogenic media. **(D-O)** Skeletal preparations of *Sp7Cre;Slc1a5^fl/+^* and *Sp7Cre;Slc1a5^fl/fl^* mice at E14.5 (n=5), E15.5 (N=8) and E16.5 (N=5). Arrows denote reduced mineralization. Isolated humeri shown in **(F)**, **(J)** and **(N).** Blue dotted lines denotes the control overall humerus length. Red dotted lines denote control mineralized area. Images quantified in **(G, K and O)**. **(P-S)** A representative skeletal preparation **(P-Q)** or **(R-S)** Representative Micro computed tomography (μCT) (N=6) used to quantify BV/TV(%) and **(T-U)** von Kossa staining on *Sp7Cre;Slc1a5^fl/+^* and *Sp7Cre;Slc1a5^fl/fl^* (N=4) knockout mice at P0. Error bars depict SD. * p≤ 0.05, ** p≤ 0.005, *** p≤ 0.0005, **** p≤ 0.00005, by an unpaired 2-tailed Student’s *t*-test.

### *Slc1a5* is required for osteoblast differentiation and proliferation

To determine how *Slc1a5* regulates bone development, we first characterized the cellular effects of *Slc1a5* ablation on osteoblasts. Because we observed a consistent skull phenotype at all stages and *Sp7Cre* is expressed in both osteoblasts and hypertrophic chondrocytes in the developing limb^44^, we focused our analyses on the skull which is formed by intramembranous ossification and does not involve a cartilaginous intermediate^45^. Von Kossa staining confirmed there was delayed mineralization in *Sp7Cre;Slc1a5^fl/fl^* calvariae at E15.5 (Fig. 2A-D). This was not due to changes in the number of osteoblast progenitors as we observed no significant difference in the number of *Sp7:GFP* expressing osteoblast progenitors per bone area in *Sp7Cre;Slc1a5^fl/fl^* mice compared to *Sp7Cre;Slc1a5^fl/+^* wild type littermates (75.3±11.6% vs 64.7±9.5% in *Sp7Cre;Slc1a5^fl/+^* and *Sp7Cre;Slc1a5^fl/f^* animals respectively) (Fig. 2E-F). However, we did observe a significant reduction in the proportion of Sp7^GFP+^ cells that were positive for proliferating cell nuclear antigen (PCNA) (25.9±4.6% vs 15.6±2.9% in *Sp7Cre;Slc1a5^fl/+^* or *Sp7Cre;Slc1a5^fl/fl^* respectively, p<0.05) (Fig. 2G-J) suggesting *Slc1a5* is required for proliferation of *Sp7* expressing osteoblast progenitors. We next evaluated osteoblast differentiation using *in situ* hybridization. We did not observe significant differences in the expression of early osteoblast genes including alkaline phosphatase (Alpl, as determined by *in situ* staining) or *Col1a1* (Fig. 2K-N). Conversely, *Sp7Cre;SIcla5^fl/fl^* animals had reduced expression of the osteoblast differentiation genes *Spp1* and *Ibsp* at both E15.5 and postnatal (P) day 0 (P0) (Fig. 2O-T and Fig. S2A-B). Similarly, the mature osteoblast gene *Bglap* was highly reduced at P0 in *Sp7Cre;Slc1a5^fl/fl^* compared to *Sp7Cre;Slc1a5^fl/+^* littermates (Fig. 2U-V). Similar results were observed in developing humeri at e15.5 (Fig. S2C-V). Consistent with these observations, primary calvarial cells isolated from *Sp7Cre;Slc1a5^fl/fl^* mice incorporated less EdU and were characterized by reduced osteoblast differentiation and matrix mineralization *in vitro* (Fig. 2W-X). This is likely a direct effect of loss of *Slc1a5* activity as acute ASCT2 inhibition using GPNA reduced calvarial cell proliferation and prevented matrix mineralization *in vitro* (Fig. S2W-Y). Collectively, these data indicate *Slc1a5* acts cell-autonomously to regulate osteoprogenitor proliferation and osteoblast differentiation.

**Fig. 2.**
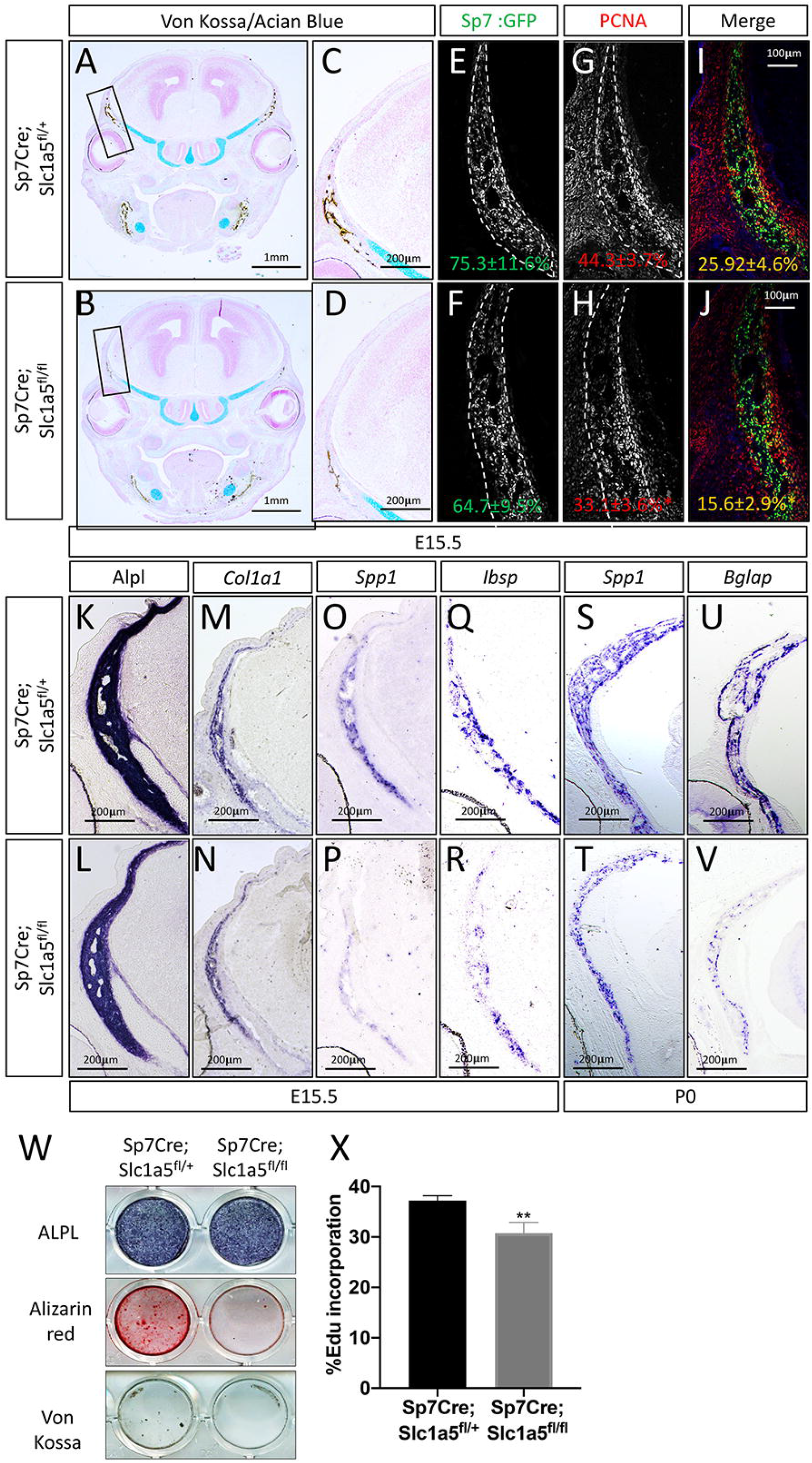
Slc1a5 is necessary for osteoblast proliferation and differentiation. **(A-D)** von Kossa/Alcian blue staining on *Sp7Cre;Slc1a5^fl/+^* **(A,C)** and *Sp7Cre;Slc1a5^fl/fl^* **(B,D)** (N=4) at E15.5. **(E-J)** Representative immunofluorescent staining for Proliferating Cell Nuclear Antigen (PCNA) used to quantify proliferation. Endogenous GFP shown in **(G-H)** used to quantify PCNA/GFP double positive cells. The numbers in each panel represent the percent GFP, PCNA or double positive cells per GFP positive bone area (dotted line). **(K-V)** Representative alkaline phosphatase (ALPL) staining **(K-L)** or *In situ* hybridization **(M-V)** for *Col1a1, Sppl, Ibsp* at E15.5 (N=4) and *Spp1, Bglap* at P0 (N=3). **(W)** Functional assays of osteoblast differentiation in cOB isolated from *Sp7Cre;Slc1a5^fl/+^* and *Sp7Cre;Slc1a5^fl/fl^* mice. cultured for 14 days in osteogenic media. **(X)** Graphical depiction of EdU incorporation in cOB cells isolated from *Sp7Cre;Slc1a5^fl/+^* and *Sp7Cre;Slc1a5^fl/fl^* mice. Error bars depict SD. * p≤ 0.05, ** p≤ 0.005. by an unpaired 2-tailed Student’s *t*-test.

### *Slc1a5* inhibition reduces protein synthesis in osteoblasts

We next sought to understand how *Slc1a5* regulates proliferation and differentiation. Because *Slc1a5* encodes a neutral amino acid transporter, we hypothesized that *Slc1a5* ablation would primarily affect protein synthesis. To test this hypothesis, we first evaluated protein synthesis directly by monitoring the incorporation of ^3^H-proline into protein. *Slc1a5* targeting significantly reduced ^3^H-proline incorporation into both collagen and total protein (Fig. S3A-B). Likewise, primary calvarial cells isolated from *Sp7Cre;Slc1a5^fl/fl^* mice had significantly reduced protein and collagen synthesis rates *in vitro* (Fig. 3A-B). These results indicate *Slc1a5* is required for robust protein synthesis in osteoblasts. To test the validity of this conclusion, we evaluated the production of osteoblast proteins *in vivo*. Consistent with the *in vitro* data, *Sp7Cre;Slc1a5^fl/fl^* mice were characterized by significantly less COL1A1 in both the calvariae and long bones at e15.5 and P0 (Fig.s 3C-H and S3C-H). It is important to note, *Col1a1* mRNA expression was not affected in these mice indicating *Slc1a5* provides amino acids required for robust COL1A1 synthesis. Similarly, *Sp7Cre;Slc1a5^fl/fl^* mice had a significant reduction in OSX protein expression as we observed a reduction in the proportion of OSX expressing cells despite no change in *Sp7^GFP^* expression (used as a proxy for *Sp7*) in *Sp7Cre;Slc1a5^fl/fl^* calvariae (Fig. 3I-N). Thus, *Slc1a5* provides amino acids to support the synthesis of proteins like OSX to regulate terminal osteoblast differentiation and COL1A1 necessary for bone matrix production.

**Fig. 3.**
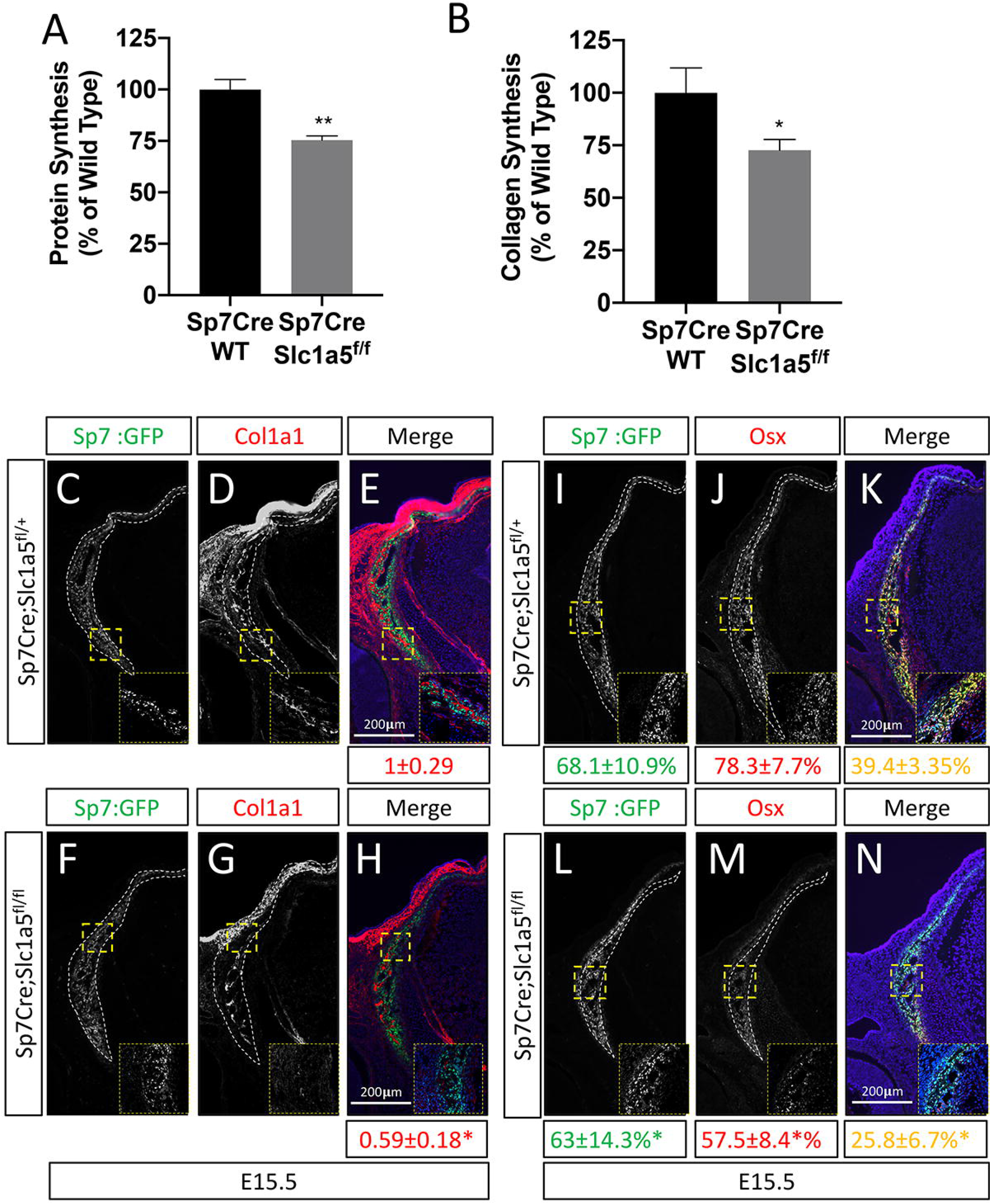
Slc1a5 is required for robust protein and matrix synthesis in osteoblasts. **(A-B)** Radiolabeled H-proline incorporation into total protein **(A)** or collagen **(B)** in cOB cells isolated from *Sp7Cre;Slc1a5^fl/+^* and *Sp7Cre;Slc1a5^fl/fl^* mice. **(C-N)** Representative immunofluorescent staining for Collagen Type 1 (COL1A1) **(C-H)** or OSX **(I-N)** at E15.5 in *Sp7Cre;Slc1a5^fl/+^* **(C,D,E,I,J,K**) and *Sp7Cre;Slc1a5^fl/fl^* **(F,G,H,L,M,N)** mice. Endogenous GFP shown in **(C,F,I,L)**. Col1a1 intensity was quantified in the GFP positive region. Endogenous GFP shown in **(I, L)** was used to quantify OSX/GFP double positive cells. The numbers in each panel represent the percent GFP, OSX or double positive cells per GFP positive bone area (dotted line). Inset images show 60x magnification of the indicated region. * p≤ 0.05, by an unpaired 2-tailed Student’s *t*-test

### *Slc1a5* provides glutamine and asparagine to regulate amino acid homeostasis

We next sought to define the molecular substrates of ASCT2 in osteoblasts. First, we determined the effect of *Slc1a5* knockout on downstream metabolites using mass spectrometry. *Slc1a5* targeting significantly diminished the intracellular abundance of many amino acids including reported substrates of ASCT2 (e.g. asparagine, glutamine and alanine) as well as amino acids not known to be transported by ASCT2 (e.g. glutamate, lysine, histidine, aspartate, glycine and proline) (Fig. 4A). Moreover, *Slc1a5* deletion also reduced the abundance of select TCA cycle intermediates including fumarate, malate, citrate and a-ketoglutarate (Fig. S4A). Interestingly, the uptake of many of these amino acids was unaffected in *Slc1a5* deficient cells as only glutamine and to a lesser extent asparagine uptake was diminished in *Slc1a5* targeted calvarial cells (Fig. 4B). Conversely, we observed a compensatory increase in the uptake of lysine in *Slc1a5* deficient calvarial cells (Fig. 4B). Similar results were obtained upon acute ASCT2 inhibition indicating *Slc1a5/ASCT2* transports glutamine and asparagine in osteoblasts (Fig. S4B). *Slc1a5* deficient cells had many cellular changes consistent with decreased amino acid concentrations. For example, *Slc1a5* inhibition significantly increased EIF2a Ser51 phosphorylation, a marker of amino acid depletion (Fig. 4C and S4C). Additionally, we observed a significant reduction in mTORC1 activity, as both ribosomal protein S6 Ser240/244 phosphorylation and Eif4ebp1 Ser65 phosphorylation were significantly reduced in *Slc1a5* deficient calvarial cells (Fig. 4C). Interestingly, mTOR activation was not affected by acute ASCT2 inhibition indicating decreased mTORC1 signaling may be a secondary effect of *Slc1a5* deletion. Collectively, these data indicate *Slc1a5* provides glutamine and asparagine to regulate intracellular amino acid homeostasis in osteoblast progenitors.

**Fig. 4.**
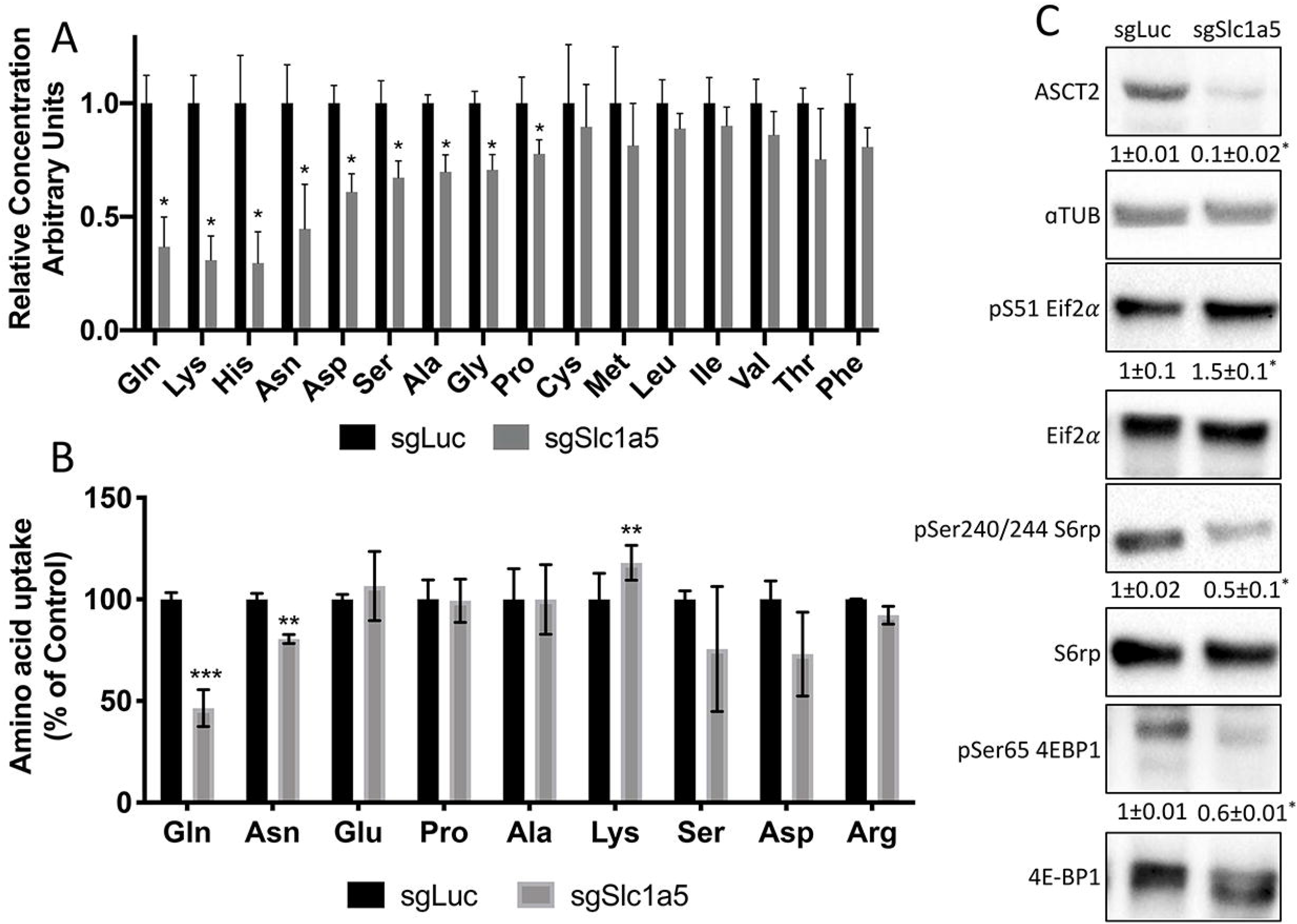
Slc1a5 provides glutamine and asparagine to maintain amino acid homeostasis. **(A)** Effect of *Slc1a5* targeting on intracellular amino acid concentration measured by mass spectrometry. **(B)** Effect of *Slc1a5* targeting on the uptake of indicated radiolabeled amino acids. **(C)** Western Blot analyses of the effect of *Slc1a5* targeting on mTORC1 signaling and Eif2a phosphorylation. Phospho-proteins normalized to respective total protein. ASCT2 normalized to α-tubulin. sgRNAs targeting luciferase were used as a negative control. Fold change ± SD for *sgSlc1a5* over *sgLuc* in 3 independent experiments. * p≤ 0.05 by an unpaired 2-tailed Student’s *t*-test

### Glutamine depletion mimics the effects of *Slc1a5* deletion

We next sought to understand the importance of both glutamine and asparagine for cellular function. To do this, we cultured naïve calvarial cells in the absence of glutamine or in media treated with asparaginase to deplete asparagine. Depletion of either glutamine or asparagine from the media similarly inhibited the induction of terminal osteoblast genes *Ibsp* and *Bglap* and prevented matrix mineralization (Fig. 1B-C and Fig. 5A-B). While this was reminiscent of *Slc1a5* ablation, it is important to note that glutamine withdrawal had a more profound effect on osteoblast differentiation compared to either asparagine depletion or *Slc1a5* ablation. We next evaluated mTORC1 activity and EIF2a phosphorylation. Depletion of glutamine, but not asparagine, mimicked the effects of *Slc1a5* targeting on both EIF2a Ser51 phosphorylation and mTORc1 activity (Fig. 5C). Consistent with this, cells cultured in the absence of glutamine had reduced COL1A1 expression and decreased proliferation like *Slc1a5* deficient cells (Fig. 5C-D). On the other hand, culturing cells in the absence of asparagine had no discernable effect on either pSer51 Eif2a, pSer240/244 S6rp or COL1A1 expression and enhanced proliferation as determined by increased EdU incorporation (Fig. 5C-D). Importantly, culturing cells in the absence of either glutamine or asparagine had no effect on cell viability (Fig. 5E). From these data we conclude *Slc1a5*/ASCT2 primarily provides glutamine to regulate amino acid homeostasis necessary for proliferation and osteoblast differentiation. Additionally, *Slc1a5*/ASCT2 provides asparagine which is essential for terminal osteoblast differentiation and matrix mineralization.

**Fig. 5.**
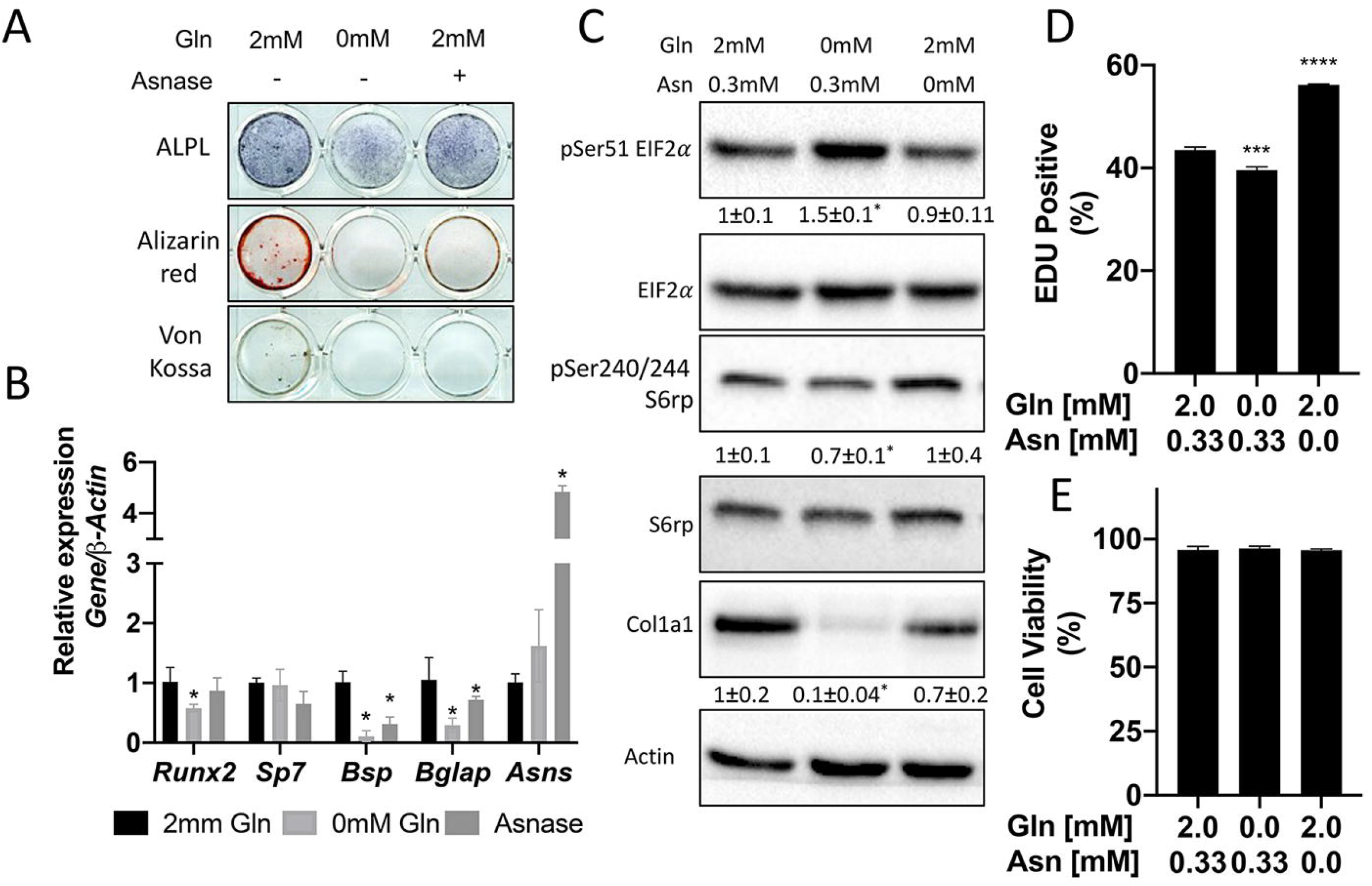
Glutamine and asparagine are required for osteoblast differentiation. **(A-B)** Functional assays **(A)** or qRT-PCR analyses **(B)** of the effect of glutamine withdrawal or asparaginase treatment on cOB cultured in osteogenic media for 14 days. **(C)** Western blot analyses of the effect of glutamine or asparagine withdrawal on mTORC1 signaling, Eif2a phosphorylation or COL1A1 expression. Phospho-proteins normalized to respective total protein. COL1A1 normalized to beta-actin. Fold change ± SD for 3 independent experiments. * p≤ 0.05 by an unpaired 2-tailed Student’s *t*-test. **(D-E)** Effect of glutamine or asparagine withdrawal on EdU incorporation **(D)** or cell viability **(E)** as determined by Annexin V staining.

### Glutamine and asparagine dependent amino acid synthesis is essential for protein synthesis

We next investigated how osteoblast utilize glutamine and asparagine. Since *Slc1a5* reduced intracellular amino acids, we hypothesized osteoblasts rely on glutamine and asparagine metabolism to maintain cellular amino acid pools. To test the validity of this hypothesis, we traced the relative contribution of glutamine or asparagine into amino acids directly. Consistent with previous reports, glutamine carbon contributes to all TCA cycle intermediates and was significantly enriched in several amino acids (e.g., glutamate, aspartate, alanine and proline) found to be reduced in *Slc1a5* targeted cells (Fig. 6A, S5A). Similarly, glutamine nitrogen was significantly enriched in glutamate, aspartate, alanine, serine, glycine and proline (Fig. 6B). Consistent with the minor effects of asparagine withdrawal on markers of amino acid depletion, asparagine carbon was enriched only in aspartate, malate, fumarate and citrate (Fig. 6A, Fig. S5A). On the other hand, asparagine nitrogen was enriched in aspartate, glutamate, proline, serine and alanine (Fig. 6A-B). Thus, glutamine contributes both carbon and nitrogen for amino acid biosynthesis whereas asparagine carbon is used only for aspartate biosynthesis while asparagine nitrogen is used in transamination reactions. Importantly, the amino acids derived from either glutamine carbon (e.g., Glu, Ala, Asp, Pro) and nitrogen (e.g., Glu, Ala, Asp, Ser, Gly and Pro) and asparagine carbon (e.g., Asp) and nitrogen (e.g., Asp, Pro, Ala, Ser) were significantly enriched in total protein (Fig. 6C-D). These data indicate *Slc1a5* provides glutamine and asparagine important for the *de novo* synthesis of many amino acids (e.g., Glu, Asp, Ala, Ser, Gly and Pro) that support nascent protein production in osteoblasts.

**Fig. 6.**
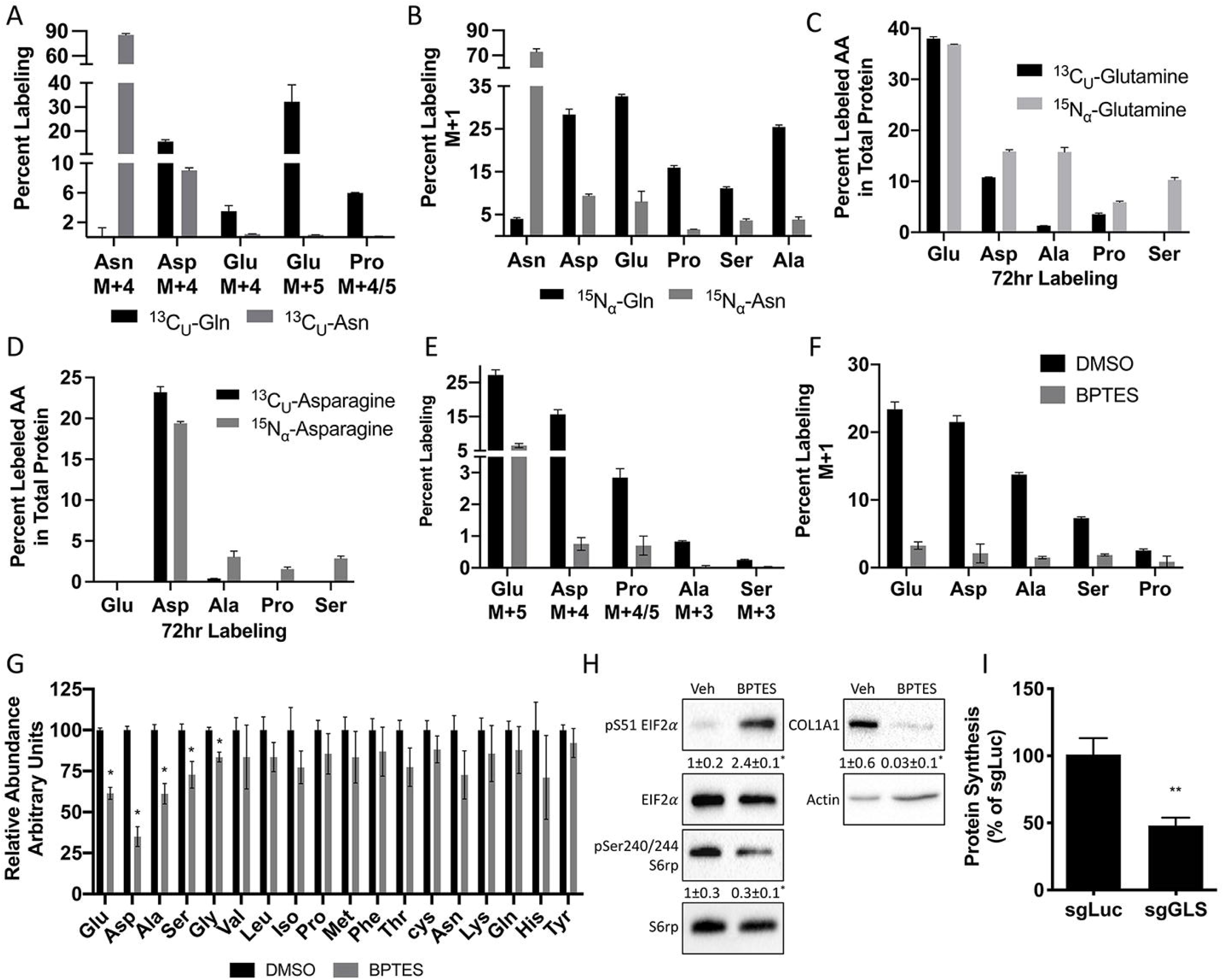
Glutamine and asparagine are utilized for de novo amino acid biosynthesis. **(A-B)** Fractional contribution of [U-^13^C]glutamine or [U-^13^C]asparagine **(A)** or [α-^15^N]glutamine or [α-^15^N]asparagine **(B)** to asparagine, aspartate, glutamate, proline, serine and alanine. **(C-D)** Fractional contribution of [U-^13^C]glutamine, [U-^13^C]asparagine, [α-^15^N]glutamine or [α-^15^N]asparagine to asparagine, aspartate, glutamate, proline, serine and alanine in total protein. **(E-F)** Effect of BPTES treatment on the fractional contribution of [U-^13^C]glutamine **(E)** or [α-^15^N]glutamine **(F)** to amino acids. **(G)** Effect of GLS inhibition on intracellular amino acid concentration measured by mass spectrometry. **(H)** Western blot analyses of the effects of BPTES treatment on mTORC1 signaling, Eif2a phosphorylation and COL1A1 expression. Phospho-proteins normalized to respective total protein. COL1A1 normalized to beta-actin. Fold change ± SD for 3 independent experiments. * p≤0.05 by an unpaired 2-tailed Student’s *t*-test. **(I)** Effect of GLS inhibition on protein synthesis as determined by the rate of ^3^H Proline incorporation into total protein. Error bars depict SD.

Next, we sought to determine if glutamine dependent amino acid synthesis was required for the high rate of protein synthesis in osteoblasts. We focused on glutamine because glutamine was more widely used for amino acid synthesis, and glutamine withdrawal more completely phenocopied the cellular effects of *Slc1a5* targeting. To do this, we inhibited the enzyme glutaminase (GLS) which catalyzes the first rate-limiting step in glutamine metabolism. GLS inhibition using BPTES resulted in cellular effects similar to *Slc1a5* ablation. GLS inhibition significantly reduced glutamine carbon and nitrogen contribution to amino acid synthesis and reduced the intracellular concentrations of glutamate, aspartate, alanine, serine, and glycine concentrations (Fig. 6E-G). Consistent with reduced amino acid concentrations, GLS inhibition increased the presence of uncharged Glu, Asp and Pro tRNA without affecting either Gln or Val tRNA charging (Fig. S5D). Finally, GLS inhibition induced eIF2α Ser51 phosphorylation and reduced S6 Ser240/244 phosphorylation (Fig. 6H). Consistent with these molecular changes, GLS inhibition significantly reduced EdU incorporation and COL1A1 expression (Fig. 6H-I). Importantly, *Gls* targeting reduced overall protein synthesis like *Slc1a5* targeting (Fig. S5E-G, 6I). These data highlight the importance of *de novo* amino acid synthesis to maintain amino acid homeostasis and promote proliferation and protein synthesis in osteoblasts.

## Discussion

Here we present data demonstrating that ASCT2 is a critical regulator of osteoblast proliferation, differentiation and bone formation. Osteoblasts increase *Slc1a5* expression as they undergo differentiation and increase bone matrix production. Genetically inhibiting *Slc1a5* activity in preosteoblasts results in delayed endochondral and intramembranous ossification. Delayed bone development is the result of decreased osteoprogenitor proliferation as well as a reduction in overall osteoblast differentiation. *Slc1a5* deficient osteoblasts have reduced protein synthetic activity which manifests in reduced OSX and COL1A1 protein expression. COL1A1 is the major organic component of bone matrix and OSX is required for terminal osteoblast differentiation^1^. Mechanistically, *Slc1a5* primarily provides glutamine and to a lesser extent asparagine in osteoblasts. Glutamine appears to be a critical nutrient at all stages of osteoblast differentiation whereas asparagine is more important for terminal osteoblast differentiation.

Proliferation is one of the initial stages of the osteoblast differentiation process. Proliferation is associated with increased demand for glucose and amino acids to support both nucleotide and non-essential amino acid (NEAA) biosynthesis required for cell division^8,10,12,46^. To fulfil this need, proliferating cancer cells increase amino acid transporter expression to enhance amino acid supply^27,38,43,47–50^. Here we find that *Slc1a5* provides glutamine and asparagine to osteoblasts, but that only glutamine is required for proliferation. This is consistent with recent reports that asparagine is important for proliferation only when glutamine is limited^51–53^. Glutamine has long been recognized as an essential nutrient in proliferating cells^54^, and recent studies found glutamine and its metabolism are critical for proliferation in osteoblast progenitor cells^21,55^. Glutamine catabolism provides amino acids and likely other intermediate metabolites to support proliferation. Consistent with this conclusion, non-essential amino acid supplementation could rescue proliferation of GLS deficient osteoblasts^55^. This is reminiscent of proliferating cancer cells which increase amino acid biosynthesis to provide carbon, nitrogen and other metabolites to support both protein and nucleotide biosynthesis as well as cellular redox homeostasis^25,56–61^.

Our data demonstrates that osteoblasts rely on a unique mechanism where they increase the supply of glutamine and asparagine which are used to maintain amino acid homeostasis. Interestingly, glutamine was more widely used for NEAA biosynthesis as our tracing experiments found glutamate, alanine, serine, aspartate, proline, and glycine were all synthesized from either glutamine carbon or nitrogen. Conversely, only aspartate was synthesized using asparagine carbon whereas Asp, Glu, Ala, and Ser were synthesized using asparagine nitrogen albeit at lower levels compared to glutamine (Fig. 6A-B). It is interesting that glutamine and asparagine metabolism impinge upon aspartate biosynthesis in osteoblasts. Aspartate synthesis is critical for proliferation by providing both carbon and nitrogen for synthetic reactions^60,61^. Thus, glutamine (and asparagine when glutamine is limiting) likely regulate proliferation in part by providing both carbon and nitrogen for aspartate biosynthesis^52^. Aspartate synthesis likely occurs as part of the malate aspartate shuttle which is critical to regenerate oxidized NAD^+^ to maintain high rates of glycolysis in osteoblasts^62^. While we don’t completely understand the metabolism of aspartate in osteoblasts, the aspartate derived from both glutamine and asparagine was enriched in total protein (Fig. 6E). Interestingly, while glutamine contributed more to aspartate biosynthesis than asparagine, asparagine-derived aspartate was more highly enriched in total protein. This suggests disparate pools of aspartate exist in osteoblasts and that the metabolic fates for these glutamine- or asparagine-derived aspartate pools vary. For example, glutamine derived aspartate may contribute more to nucleotide biosynthesis to facilitate osteoblast proliferation whereas asparagine derived aspartate is used more for protein and matrix synthesis. Future studies are warranted to investigate this possibility.

During differentiation, osteoblasts increase their capacity for protein synthesis and secretion. Here we find that *Slc1a5* expression is required for robust protein synthesis in osteoblasts. This effect is likely regulated by multiple factors. First, *Slc1a5* provides glutamine and asparagine that are likely directly incorporated into nascent protein. Second, our tracing data shows both glutamine and asparagine are used for *de novo* amino acid synthesis to support protein synthesis. Third, *Slc1a5* provides glutamine to activate the mechanistic target of rapamycin complex 1 (mTORC1), an important regulator of both osteoblast proliferation and differentiation^63–67^. mTORC1 is tightly regulated by amino acid availability and functions to increase protein synthesis when amino acids are available. Consistent with this, *Slc1a5* deficient osteoblasts had decreased mTORC1 activation and reduced protein synthesis (Fig.s 3A-B and 4C). Finally, *Slc1a5* deficient osteoblasts were characterized by robust activation of the integrated stress response (ISR). When amino acids are limiting, uncharged tRNA activates the kinase general control nonderepressible 2 (GCN2) which phosphorylates eukaryotic initiation factor 2a (eIF2a) to attenuate global protein synthesis^68–71^. These molecular changes in *Slc1a5* deficient osteoblasts are primarily the result of reduced glutamine uptake as our drop out experiments found glutamine but not asparagine withdrawal reduced mTORC1 activation and increased ISR like *Slc1a5* ablation (Fig. 5C). Moreover, inhibiting glutamine catabolism similarly affected mTORC1 and ISR activation (Fig. 6H). Thus, *Slc1a5* provides glutamine used to maintain amino acid homeostasis to regulate protein synthesis directly and indirectly downstream of mTORC1 and GCN2 dependent ISR.

## Methods

### Mouse Strains

*C57B1/6J*(RRID:IMSR_JAX:000664), *Rosa26Cas9* (RRID:1MSR_JAX:024858), *Sp7tTA;tetOeGFP/Cre* (RRID:IMSR_JAX:006361), and *Rosa26Flpe* (RRID:1MSR_JAX: 003946) mouse strains were obtained from the Jackson Laboratory. The *Slc1a5^flox^* mouse strain was generated by the Duke Cancer Institute Transgenic Core facility. Briefly, LoxP sites flanking exon 2 and a frt PGK neo-cassette were inserted in to the endogenous *Slc1a5* locus using homologous recombination (Fig. S1). The chimeric mice containing the targeting vector were then crossed to the *Rosa26Flpe* mouse to remove the neomycin cassette. All mice were housed at 23°C on a 12-hour light /dark cycle and maintained on PicoLab Rodent Diet 290 (Lab diet#5053, St. Louis, MO). Timed pregnant females euthanized, and embryos were analyzed at E14.5, E15.5, E16.5 and P0. All animal studies were approved by the animal studies committees

### Cell culture

P3 pups were euthanized and the parietal and frontal bones were isolated according to standard protocols. The membranous tissue was removed, and the bones were washed and cleaned with PBS. The calvaria was chopped and four sequential 10 min digestions were performed with 1mg/ml Collagenase P in a shaking incubator at 37°C. The first digestion was discarded, and the rest were combined. The digested calvarial cells were centrifuged and plated with α-MEM (GIBCO) supplemented with 15% FBS (Invitrogen). The cells were seeded at 50,000 cells/ml. To initiate osteoblast differentiation the growth media was replaced with α-MEM with 10% FBS, 50 mg/ml ascorbic acid (Sigma) and 10 mM β-glycerophosphate (Sigma). RNA was isolated at day 7 after mineralization media was added. At day 14, alizarin red and Von kossa staining was performed to visualize matrix mineralization. Alkaline phosphatase staining was performed using the one-step nitro-blue tetrazolium (NBT) and 5-bromo-4-chloro-3’-indolyphosphate p-toluidine salt (BCIP) solution (Thermofisher).

### CRISPR/Cas9 targeting

sgRNA vectors were purchased from the Genome Engineering and iPSC Center at Washington University School of Medicine. cOB were isolated from *Rosa^Cas9^* mice and were infected with five lentiviral delivered short guide RNAs targeting exons 4-6 of either *Slc1a5* (sgSlc1a5) or *Gls* (sgGls) (Fig. S1 and S6). As a control cOB were infected with sgRNAs targeting the open reading frames of Luciferase and mCherry as a control (designated sgLuc in text and Fig.s). sgRNA sequences are listed in Table 2. To make virus, 293T cells were cotransfected with the lentiviral vector expressing short guide RNAs, pMD2.g and psPax2. After 48h of transfection the media containing the virus was collected and filtered through 0.45 *μ*m filter. cOB were cultured to 50% confluency and were infected for 24 hours followed by recovery in regular media for another 24 hours.

**Table 1.**
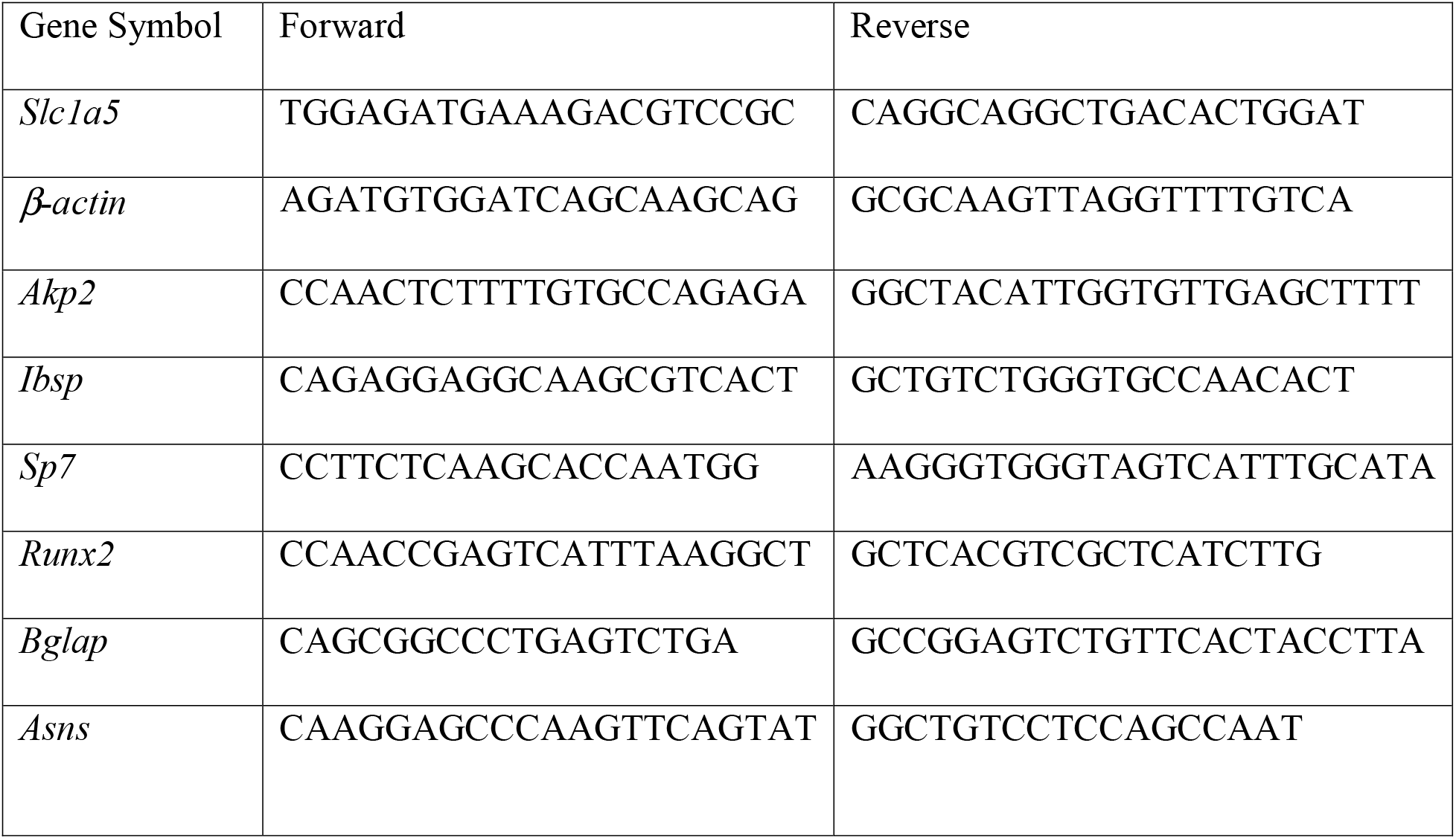
RT-PCR primer sequences.

**Table 2.**
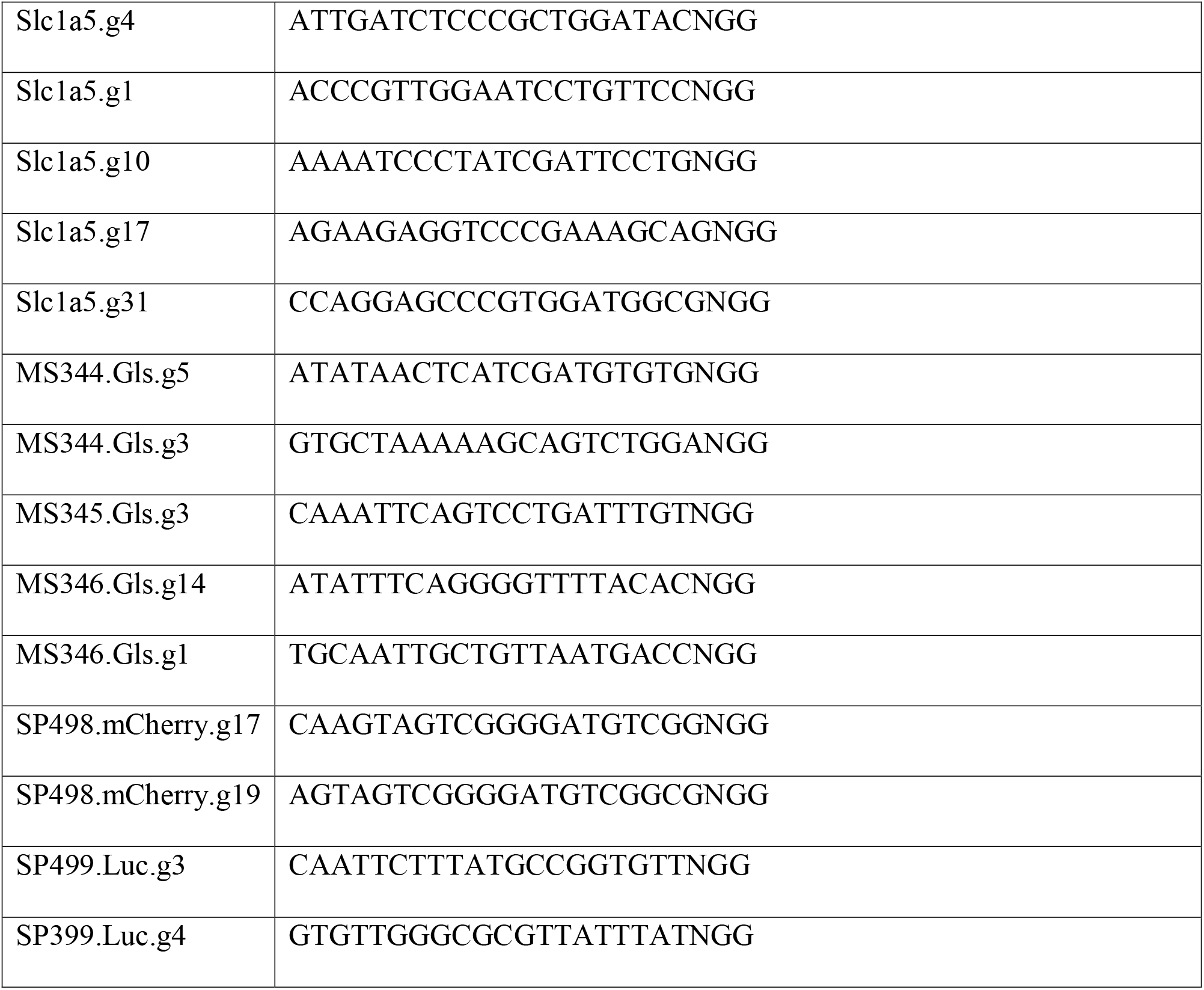
sgRNA sequences.

### Uptake Assays and Proline incorporation assay

Confluent primary cells were washed ones with PBS and two additionally washes with Krebs Ringer Hepes (KRH) (120mM NaCl, 5mM KCl, 2mM CaCl_2_, 1mM MgCl_2_, 25mM NaHCO_3_, 5mM HEPES, 1mM D-Glucose, pH 8). Cells were then treated for 5 min with KRH containing 4 μCi/mL of either L-(2,3,4-^3^H)Glutamine, L-(^14^C)Alanine, L-(2,3,-^3^H)Proline, L-(2,3,-^3^H)Aspartic acid, L-(4,5-^3^H(N))Lysine, or L-(2,3,-^3^H)Asparagine. After 5 minutes, the uptake was terminated by washing the cells 3 times with ice cold KRH. Cells were then scraped in 1mL of dH_2_O and lysed by sonication for 1 minute with 1 second pulses at 20% amplitude. The lysate was then centrifuged and mixed with 8mL scintillation liquid. The counts per minute (CPM) were measured using a Beckman LS6500 scintillation counter. Proline incorporation assay was performed on confluent cells in a 12 well plate. The cells were washed ones with PBS and two additional washes with KRH. The cells were incubated with 4 μCi/mL L (2,3,-^3^H) Proline in KRH at 37C for 3h. The uptake was terminated with three washes of cold KRH, and the cells were scraped with 150 μl RIPA. The lysates were then cleared and 90μl of the protein was precipitated with 10% tricholoroacetic acid (TCA). The pellet was washed 3 times with 5%TCA and resuspended with 1ml 0.2N NaOH. 100μl of the lysate was saved to measure proline incorporation in total protein. The remaining lysate was separated in to two 400μl parts, to one-part 200μl of 60μm HEPES containing 10 units of collagenase was added and to the second part only HEPES was added. The lysates were incubated at 37C for two hours, followed by precipitation of undigested protein with 10%TCA. The supernatant was added to 8ml of scintillation liquid to measure proline incorporation in collagen using a Beckman LS6500 scintillation counter.

### RT-PCR

RNA was isolated using the Qiagen RNAeasy kit. 500ng of RNA was reverse transcribed to cDNA using Iscript Reverse transcription kit (Biorad). The cDNA was diluted 1:10. qPCR reaction was setup using SYBR green (Biorad), 2μl of diluted cDNA and 0.1 μM primers in technical and biological triplicates. ABI Quantstudio 3 was used to run the qPCR. The PCR cycle were 95°C for 3 min followed by 35 cycles of 95°C for 10s and 60°C for 30s. Gene expression was normalized to *ActB* followed by calculating relative expression using the 2^-(ΔΔCt)^ method. The list of primers is found in table 1.

### Western blotting

To isolate protein, cells were scraped, or bones were pulverized in RIPA lysis buffer (50 mM Tris (pH 7.4), 15 mM NaCl, 0.5% NP-40) containing protease and phosphatase inhibitors (Roche). Protein concentration was measured using the BCA method and 10μg of the protein was run on 10% polyacrylamide gel. The protein was then transferred to a nitrocellulose membrane which was blocked for 1 hour at room temperature with 5% milk in TBST (TBS, 0.1% Tween20). The blots were then incubated with primary antibodies raised against ASCT2 (RRID:AB_10621427, 1:1000), Eif2α (RRID:AB_10692650), pSer51 Eif2α (RRID:AB_2096481) pSer240/244 S6rp (RRID:AB_331682, 1:1000), S6rp (RRID:AB_331355, 1:1000), α-tubulin (RRID: AB_2619646, 1:1000) or β-actin (RRID: AB_330288, 1:2000) overnight at 4C. On day 2, the membranes were washed three times 5 min each with TBST and incubated with appropriate secondary antibody HRP goat anti-rabbit (RRID:AB_2099233) or HRP anti-mouse (RRID:AB_330924) in 5% milk in TBST for 1 hour at room temperature. The membranes were again washed three times with TBST and developed using clarity ECL substrate (Biorad) or super signal West Femto ECL.

### EdU incorporation and Annexin V assay

5000 cells/well were seeded in a 96 well plate for 12-16 hours after which the cells were incubated with 10μM EdU (5-ethynyl-2’-deoxyuridine) for 6 hours. EdU incorporation was performed using the instructions provided in the Click-iT™ EdU Cell Proliferation Imaging Kit (Invitrogen, C10337). EdU incorporation was also analyzed using Click-iT™ EdU Alexa Fluor™ 488 Flow Cytometry Assay Kit (Invitrogen, C10420) after 24h incubation with 10μM EdU. The cells were then trypsinized, permeabilized and stained as per kit instructions. Cell viability assay was performed using the Apoptosis Assay Kit (Cat# 22837).

### Skeletal prep

Timed pregnant females were euthanized, and pups were harvested at E14.5, E15.5, and P0. The embryos were skinned, eviscerated and dehydrated in 95% ethanol overnight. The mice were transferred to acetone for another night to dissolve fat tissue. After which, the tissue was stained with Alcian blue 8GX (0.03%, m/v in 70% ethanol) and Alizarin red S (0.005%, m/v in dH_2_O) solution containing 10% acetic acid and 80% ethanol. The stained skeletons were cleared in 1% KOH followed by a gradient of glycerol.

### Histology, immunofluorescence and In situ hybridization

Limbs and skulls from E15.5, E16.5 and P0 were harvested skinned and fixed overnight in 4% PFA (paraformaldehyde). Limbs from E16.5 and P0 were decalcified in 14% EDTA overnight, followed by transferring them to 30% sucrose for frozen embedding in OCT and 70% ethanol for paraffin embedding. Paraffin embedded blocks were sectioned at 5μM thickness and utilized for Alcian blue and picro-sirius, von Kossa and alcian blue using standard protocols.

Immunofluorescence was performed on 10μM frozen sections brought to room temperature and incubated with 3% H_2_O_2_ for 10min. The sections were then incubated with 1.5% goat serum in PBST (PBS with 0.1% Tween20) at room temperature for 1hour. The sections were then incubated overnight in primary antibodies COL1A1 (RRID:AB_638601,1:200), OSX (RRID:AB_2194492 1:500,), PCNA (RRID:AB_2160343,1:500) diluted in 1.5% goat serum in PBST. On day 2, the sections were washed 3 times with PBST for 5 min each and incubated with goat anti mouse 568 (RRID:AB_141359, 1:1000) for COL1A1 and goat anti Rabbit 568 (RRID:AB_143157, 1:1000) for OSX and PCNA for 30 min at room temperature. The sections were then mounted with DAPI and imaged. COL1A1 intensity was measured in the GFP positive region and OSX, GFP copositive cells were counted using Image J.

In situ hybridization was performed on 10μM frozen sections. The sections were fixed with 4% PFA for 10min at room temperature followed by 10min in acetylation solution (1.3% triethanolamine, 0.175% HCl, 0.375% acetic anhydride in dH_2_O). The sections were then washed and incubated with hybridization buffer (50% formamide (deionized), 5X SSC, pH 4.5 (use citric acid to pH), 1% SDS, 50μg/mL yeast tRNA, 50μg/mL heparin) for 2 hours in a humidified chamber. The excess hybridization buffer was removed and prewarmed probe diluted 1:10 was added to the slides, covered with parafilm and incubated at 60C overnight. The slides were immersed in 5X SSC to remove the parafilm and washed twice with 0.2X SSC for 30 min at 60 C. After an additional wash at room temperature with 0.2X SSC the slides were transferred to NTT (0.15M NaCl, 0.1M tris-Cl pH 7.5, 0.1% tween 20) for 10 min at room temperature. The slides were blocked with blocking buffer (5% heat inactivated sheep serum, 2% blocking reagent/NTT) for 2hours, followed by incubation with anti-Dig AP antibody diluted at 1:4000 in the blocking buffer overnight at 4C. On the third day, the slides were washed with NTT on rotator 3 times for 30 min each, followed by 3 5min washes with NTTML (0.15M NaCl, 0.1M tris pH 9.5, 50mM MgCl_2_,2mM Levamisole, 0.1% tween 20). The slides were then incubated with prewarmed AP substrate BM purple (ROCHE) at 37 C and monitored for desired staining. After staining was achieved the slides were rinsed in PBS 3 times for 5 min each. The slides were fixed in 0.2% glutaraldehyde in 4% PFA overnight, the slides were then mounted with glycergel and imaged.

### Micro computed tomography (uCT)

VivaCT 80, Scanco Medical AG was used for three-dimensional reconstruction, and quantification of bone parameters (threshold set at 280).

### Mass spectrometry and metabolic tracing

cOB isolated from RosaCas9 homozygous P3 pups were cultured in 6cm plates and treated as indicated. Cells were incubated with 2mM [U-^13^C]glutamine, 2mM [α-^15^N]glutamine, 0.33mM [U-^13^C] Asparagine and 0.33mM [α-^15^N] Asparagine for either 24 or 72 hours as indicated.

Cells treated for 24 hours were washed with cold PBS and isolated three times with −80°C methanol on dry ice in Eppendorf tubes containing 20nM norvaline, which is an internal standard. The extracts were vortexed and centrifuged at 12000RPM for 15min. The supernatant was completely dried with N_2_ gas. The dried residue was resuspended in 25μl methoxylamine hydrochloride (2% MOX in pyridine) and incubated for 90 minutes at 40°C on a heat block. After the incubation the samples were spun at maximum speed for 2min and 35μl of MTBSTFA+1% TBDMS was added followed by a 30-minute incubation at 60°C. The samples were centrifuged at 12000RPM for 5 minutes and the supernatant was transferred to GC vials for GC-MS analysis. For the experiments tracing labeled amino acids into protein, cells were incubated with 2mM [U-^13^C]glutamine, 2mM [α-^15^N]glutamine, 0.33mM [U-^13^C]Asparagine or 0.33mM [α-^15^N] Asparagine for 72 hours. Cells were isolated in 1ml 1M PCA and centrifuged for 10 min to precipitate the protein. The precipitate was washed twice with 1ml 70% ethanol after which 20nM norvaline was added to the samples. The protein was hydrolyzed with 1ml 6M HCl at 110°C for 18 hours. The samples were cooled and 1ml chloroform was added and vortexed to remove hydrophobic metabolites. The isolates were centrifuged for 10 minutes and 100μl of the supernatant was dried with N_2_ gas until dry, 50μl of MTBSTFA+1% TBDMS was added followed by a 30-minute incubation at 60°C. The samples were transferred to GC vials for GC-MS analysis.

### tRNA aminoacylation assay

The method is adapted from two recent reports^72^. Purified RNA was resuspended in 30mM NaOAc/HOAc (pH 4.5). RNA was divided into two parts (2*μ*g each): one was oxidized with 50mM NaIO_4_ in 100mM NaOAc/HOAc (pH 4.5) and the other was treated with 50mM NaCl in NaOAc/HOAc (pH 4.5) for 15 min at room temperature. Samples were quenched with 100mM glucose for 5min at room temperature, followed by desaltation using G50 columns and precipitation using ethanol. tRNA was then deacylated in 50mM Tris-HCl (pH 9) for 30min at 37°C, followed by another ethanol precipitation. RNA (400ng) was then ligated the 3’adaptor (5’-/5rApp/TGGAATTCTCGGGTGCCAAGG/3ddC/-3’) using T4 RNA ligase 2(NEB) for 4 h at 37°C. 1*μ*g RNA was then reverse transcribed using SuperScript III first strand synthesis system with the primer (GCCTTGGCACCCGAGAATTCCA) following the manufacturer’s instruction.

### Quantification and Statistical Analysis

All statistics were performed in Graphpad 6 software. In cell culture studies, statistical significance was determined by an unpaired 2-tailed Student’s *t*-test or one-way Anova. For uCT statistical significance was determine by a paired 2-tailed Student’s *t*-test comparing paired littermate controls. All quantifications are represented as mean ± standard deviation. A P value of less than 0.05 is considered statistically significant. All experiments were performed with n≥3 biological replicates. The sample size and statistical analysis are noted in the Figure legends.

## Acknowledgements

The authors thank Drs. Vishal Patel and Guoli Hu for critical comments on this manuscript. This work was supported by National Institute of Health R01 grants (AR076325 and AR071967) to C.M.K.

## Author Contributions

Conceptualization, C.M.K.; Investigation, D.S., Y.Y., L.S., G.Z., and C.M.K.; Writing – Original Draft, C.M.K.; Writing – Review & Editing, D.S., Y.Y., L.S., G.Z., and C.M.K.; Supervision, C.M.K.

## Declaration of Interests

The authors declare no competing interests.

## Supplemental Figure Legends

**Supplemental Fig. 1: Related to Fig. 1**.

**(A)** *Slc1a5* Crispr targeting strategy. **(B)** Western Blot analyses of the effect of *Slc1a5* targeting ASCT2 normalized to α-tubulin. sgRNAs targeting luciferase and mCherry were used as a negative control. Fold change ± SD for *sgSlc1a5* over *sgLuc* in 3 independent experiments. * p≤ 0.05 by an unpaired 2-tailed Student’s *t*-test. **(C)** *Slc1a5^fl^* targeting strategy. **(D)** Western Blot analyses of ASCT2 expression in protein isolated from Muscle, Bone and Bone marrow from *Sp7Cre;Slc1a5^fl/+^* and *Sp7Cre;Slc1a5^fl/fl^* mice. **(E-X)** Skeletal preparations of *Sp7Cre;Slc1a5^fl/+^* and *Sp7Cre;Slc1a5^fl/fl^* hindlimbs at E14.5 (n=5), E15.5 (N=8), E16.5 (N=5) and (p0). Images quantified in **(G, H, K,L,O, P,U and X)**.

**Supplemental Fig. 2: Related to Fig. 2.**

**(A-V)** In situ hybridization for *Ibsp* on p0 skull **(A-B)**, representative alkaline phosphatase (ALPL) staining **(C-D)**, von Kossa/Alcian blue staining **(E-F)**, In situ hybridization **(G-L, O-P, S-V)** for *ColX, MMP13, SP7, Col1a1, Spp1, Ibsp* at E15.5 (N=4). Representative immunofluorescent staining for OSX **(M-N)** or Collagen Type 1 (COL1A1) **(Q-R)** at E15.5 in *Sp7Cre;Slc1a5^fl/+^* and *Sp7Cre;Slc1a5^fl/fl^* mice. **(W-Y)** EdU incorporation **(W-X)**, functional assay **(Y)** of effect of GPNA (0.3mM) treatment. Error bars depict SD. * p≤ 0.05, ** p≤ 0.005. by an unpaired 2-tailed Student’s *t*-test.

**Supplemental Fig. 3: Related to Fig. 3.**

**(A-B)** Radiolabeled ^3^H-proline incorporation into total protein **(A)** or collagen **(B)** in cOB cells. **(C-H)** Representative immunofluorescent staining for Collagen Type 1 (COL1A1) at P0 in *Sp7Cre;Slc1a5^fl/+^ **(C,E,G)*** and *Sp7Cre;Slc1a5^fl/fl^* **(D,F,H)** mice. Endogenous GFP shown in **(C-D)**. The value below the merged image is the quantification of Col1a1 intensity in the GFP positive area measured from 4 mice. Inset images are 60x magnification of the indicated region. * p≤ 0.05, by an unpaired 2-tailed Student’s *t*-test.

**Supplemental Fig. 4: Related to Fig. 4.**

**(A)** Effect of *Slc1a5* targeting on intracellular concentration of Pyruvate, Lactate, Succinate, Malate, citrate, and α ketoglutarate (αKG).

**(B)** Effect of GPNA treatment on the uptake of indicated radiolabeled amino acids.

**(C)** Western Blot analyses of the effect of GPNA treatment on Eif2a phosphorylation and mTORC1 signaling. Phospho-proteins normalized to respective total protein. HCL treated cells were used as negative control. Fold change ± SD for HCL over GPNA treated in 3 independent experiments. * p≤ 0.05 by an unpaired 2-tailed Student’s *t*-test. Error bars depict SD. * p≤ 0.05, ** p≤ 0.005. by an unpaired 2-tailed Student’s *t*-test.

**Supplemental Fig. 5: Related to Fig. 6.**

**(A)** Fractional contribution of [U-^13^C]glutamine or [U-^13^C]asparagine to fumarate, malate, succinate and Citrate. **(B)** Effect of BPTES treatment on the fractional contribution of [U-^13^C]glutamine to fumarate, malate, succinate and Citrate. **(C)** Effect of BPTES on EdU incorporation. **(D)** Effect of GLS inhibition on tRNA charging. Error bars depict SD. **(E)** *Gls* Crispr targeting strategy. **(F)** PCR analysis of targeted region. Arrow denotes untargeted *Gls* band. **(G)** Western Blot analyses of the effect of *Gls* targeting. sgRNAs targeting luciferase and mCherry (denoted sgLuc) were used as a negative control.

## Notes

### Competing Interest Statement

The authors have declared no competing interest.

