## Supplementary figures and images for "*SLC1A5* provides glutamine and asparagine necessary for bone development in mice"

### Supplemental Figure 1

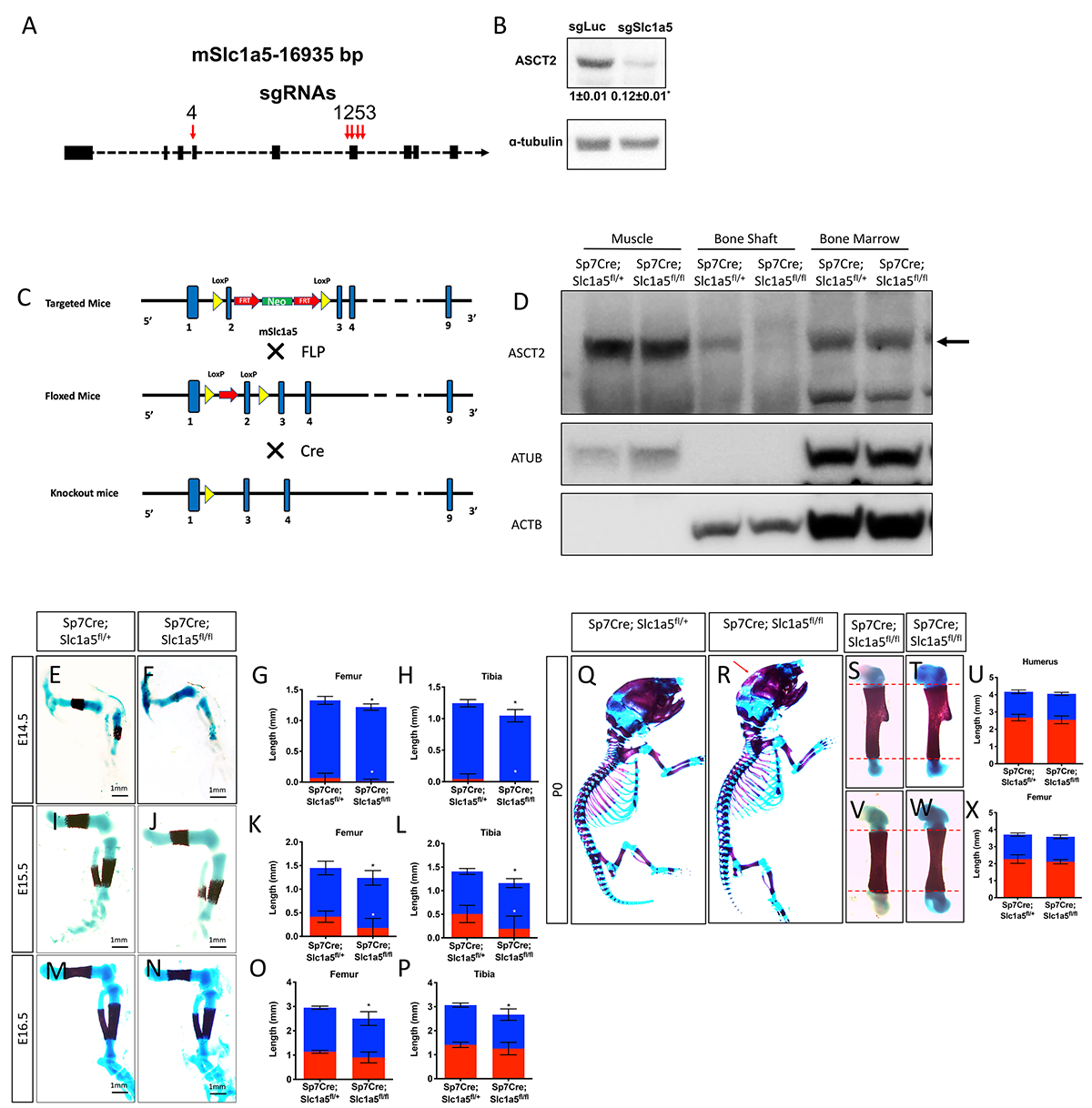

### Supplemental Figure 2

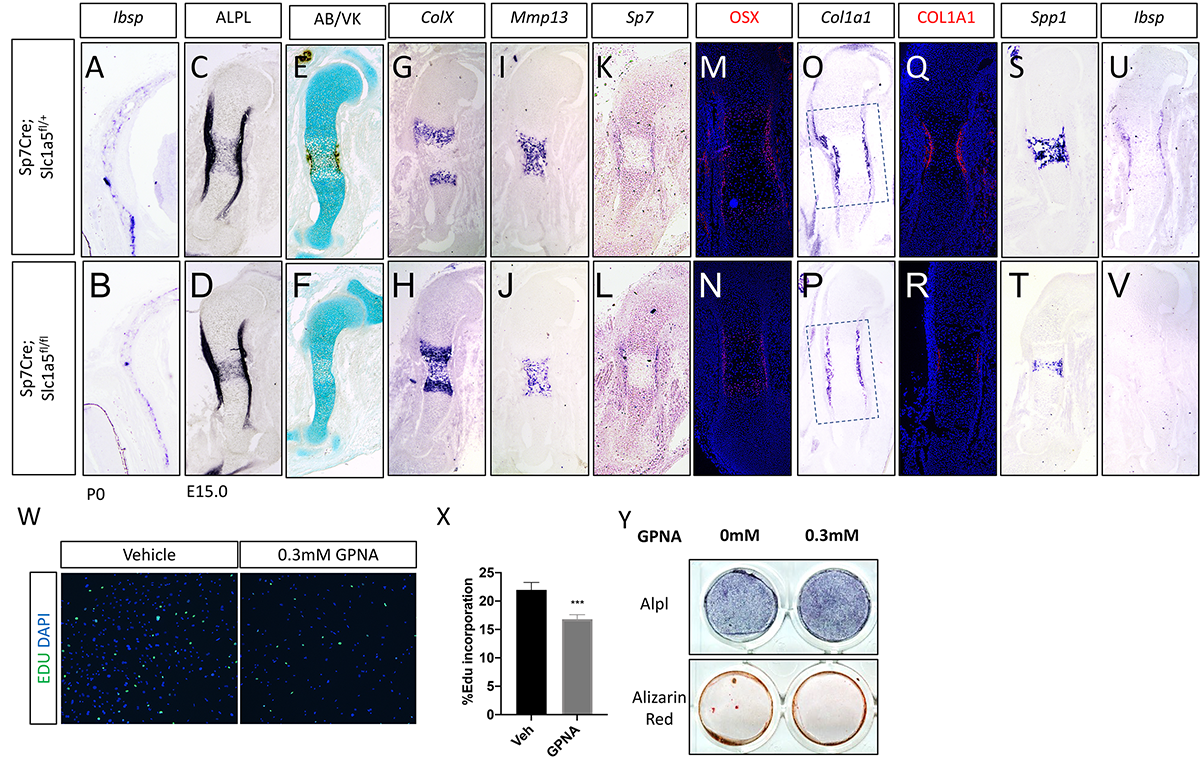

### Supplemental Figure 3

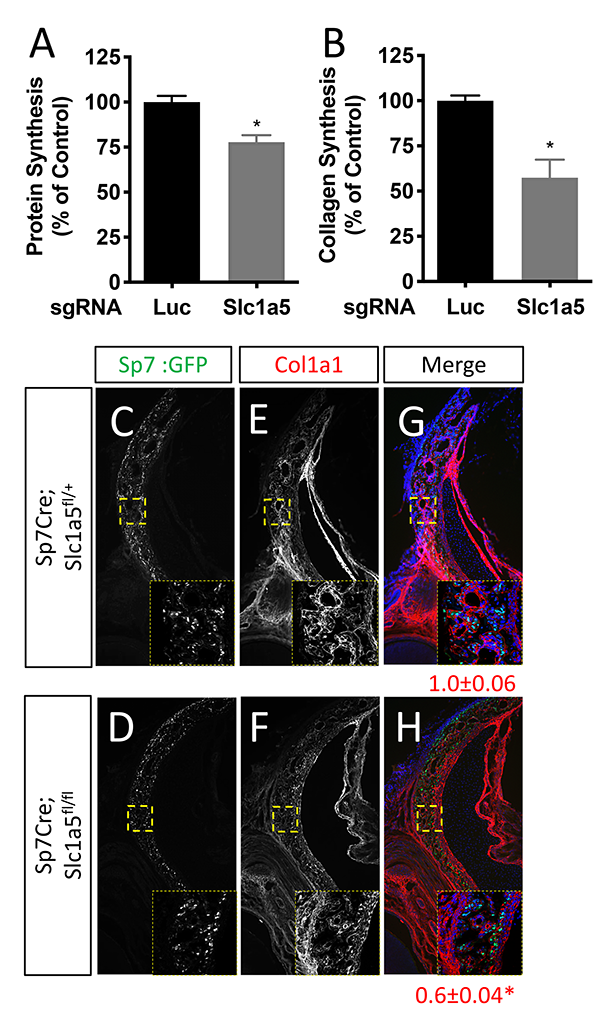

### Supplemental Figure 4

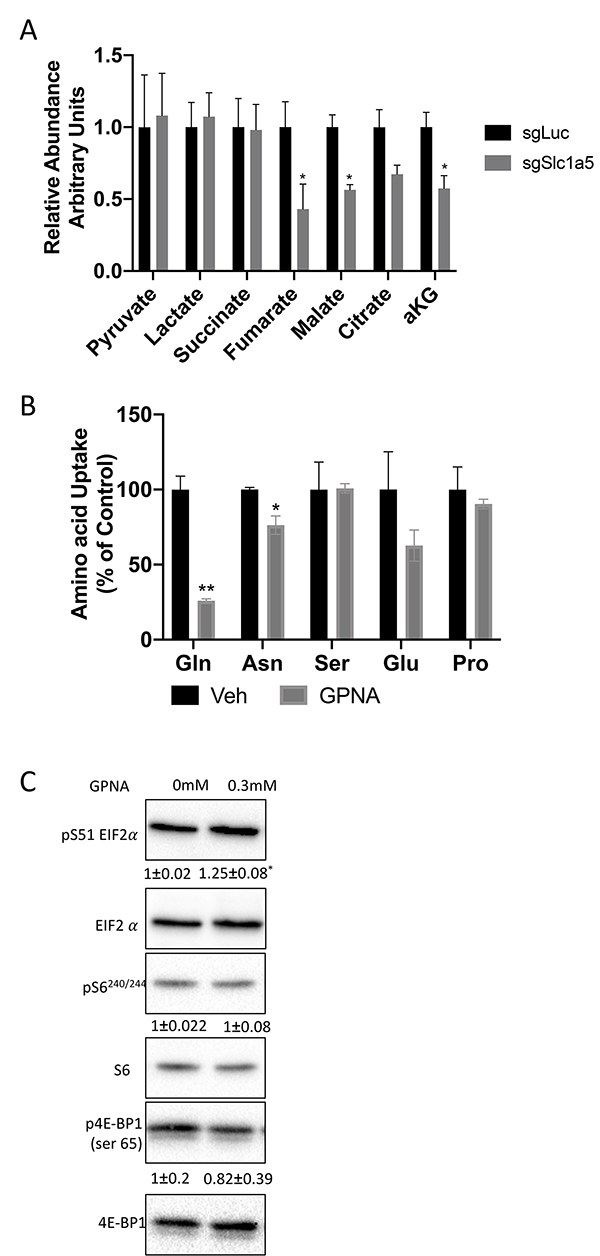

### Supplemental Figure 5

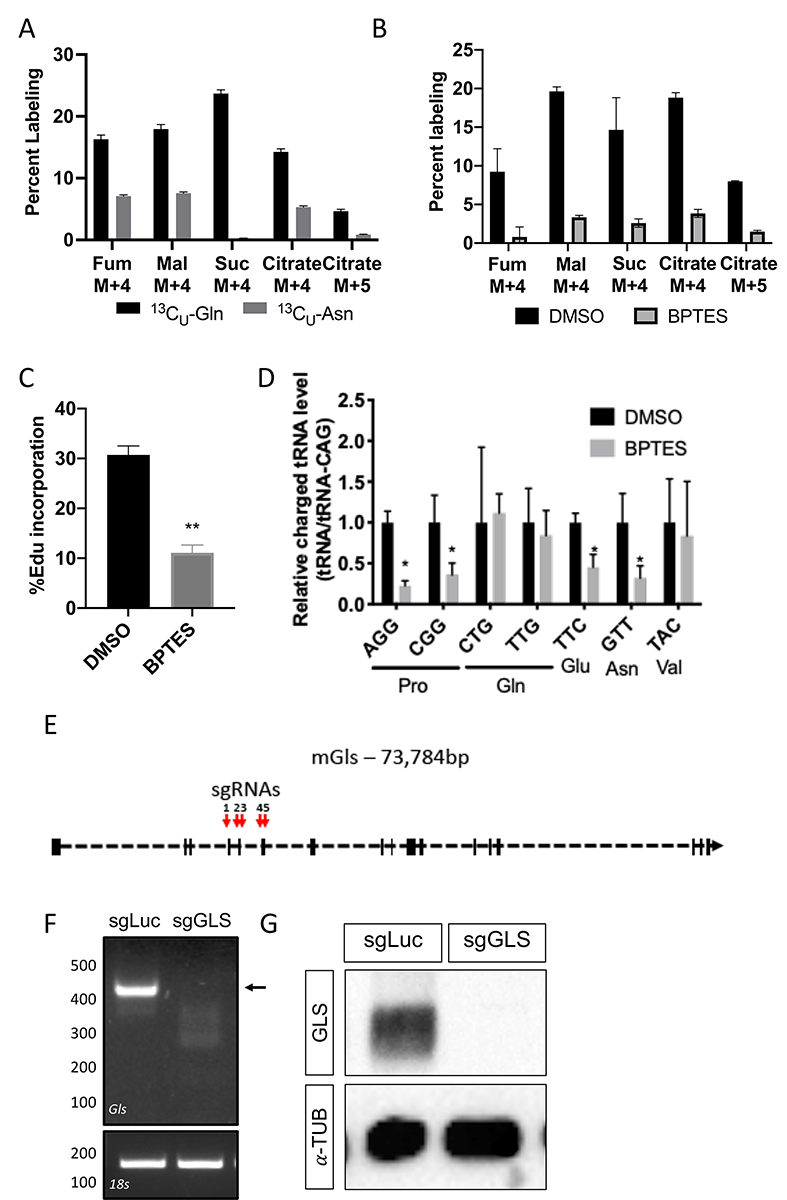
